# Gallery Game: Smartphone-based Assessment of Long-Term Memory in Adults at Risk of Alzheimer’s Disease

**DOI:** 10.1101/599175

**Authors:** Claire Lancaster, Ivan Koychev, Jasmine Blane, Amy Chinner, Christopher Chatham, Kirsten Taylor, Chris Hinds

## Abstract

Gallery Game, deployed within the Mezurio smartphone app, targets the processes of episodic memory first vulnerable to neurofibrillary tau-related degeneration in Alzheimer’s Disease, prioritising both perirhinal and entorhinal cortex/hippocampal demands. Thirty-five healthy adults (aged 40-59 years), biased towards those at elevated familial risk of dementia, completed daily Gallery Game tasks for a month. Assessments consisted of cross-modal paired-associate learning, with subsequent tests of recognition and recall following delays ranging from one to 13 days. There was a non-linear decline in memory retention with increasing delays between learning and test, with significant forgetting first reported following delays of three and five days for paired-associate recall and recognition respectively, supporting the need for ecologically valid measures of longer-term memory. Gallery Game outcomes correlated as expected with established neuropsychological memory assessments, confirming the validity of this digital assessment of episodic memory. In addition, there was preliminary support for utilising the perirhinal-dependent pattern of semantic errors during object recognition as a marker of early impairment, justifying ongoing validation against traditional biomarkers of Alzheimer’s disease.

## 1. Introduction

Detecting Alzheimer’s disease (AD) in the very earliest ‘preclinical’ stage is critical for the development of therapeutics to prevent or slow neurodegeneration. Recruitment to clinical trials targeting the initial build-up of AD-pathology, however, is limited by an over-reliance on costly, invasive biomarker screens with restricted availability (Cummings et al., 2016). Cognitive markers provide an alternative (Glymour et al., 2018), but require in-clinic assessments with trained neuropsychologists in testing sessions limited to a few hours, thus preventing the measurement of longer term memory retrieval which is pervasive in everyday life. There is therefore an urgent need for reliable, valid cognitive tools which can be deployed in individuals’ daily lives over the span of days and weeks. Digital tools facilitate high-frequency, cognitive assessment at scale (Doraiswamy, Narayan, & Manji, 2018; Harvey et al., 2017; Laske et al., 2015). Here, the utility of Gallery Game, a novel smartphone-based task, is explored in mid-age adults with a known familial bias towards late-life dementia risk.

Episodic memory is central to the design of Gallery Game, given the sensitivity of this cognitive domain to incipient AD up to 12 years prior to an official diagnosis (Amieva et al., 2008; Grober et al., 2008; Mistridis, Krumm, Monsch, Berres & Taylor, 2015). A key consideration here was the standard 30-minute retention interval (RI) used in classic episodic memory assessments (Lezak, Howieson, & Loring, 2012). By design, Gallery Game assesses much longer RIs – exceeding a week in duration – under the hypothesis that forgetting over these longer delays may be a particularly valuable marker for discriminating the prodromal and suspected preclinical stages of the disease from healthy aging (e.g., Carlesimo, Sabbadini, Fadda, & Caltagirone, 1995; Manes, Serrano, Calcagno, Cardozo, & Hodges, 2008; Walsh et al., 2014; Geurts, van der Werf, & Kessels, 2015; Reiman, 2018). This prediction has emerging support, for example the observation that *Apolipoprotein* (*APOE*) ε4, the leading genetic risk factor for late-onset AD, exerts a gene-dose effect on verbal recall and recognition in mid-age when assessed with a 7-day but not 30-minute RI (Zimmermann & Butler, 2018). Longer-term forgetting may also be particularly meaningful for patients, given that the memory performance of individuals self-referring to a memory clinic with subjective cognitive complaints can be seen to differ from that of individuals without cognitive complaints after a 7-day, but not 30-minute RI (van der Werf, Geurts, & de Werd, 2016). The assessment of longer-term RIs may therefore convey meaningful prognostic information. The burden of repeated study visits has to date limited the examination of such long-term RIs using traditional approaches (Fisher & Radvansky, 2018), however Gallery Game’ exploits the ease of mobile data collection to probe both recognition and recall across an array of long-term RIs.

The relationship between episodic memory decline to the staging of Braak pathology, specifically the initial, sequential aggregation of phosphorylated neurofibrillary tau in the perirhinal cortex (PRc), entorhinal cortex (ERc) and hippocampal regions of the anterior medial temporal lobe (aMTL) (Braak & Braak, 1991; Hirni, Kivisaari, Monsch, & Taylor, 2013; Krumm et al., 2016), was a second key consideration in the design of Gallery Game. The PRc occupies the functional apex of the ventral occipito-temporal visual processing stream, supporting the perception and representation of objects in semantic memory (Bussey, Saksida, & Murray, 2005). Hence, in line with PRc being the earliest cortical site of tau accumulation in AD, tasks loading on this ability should provide the first signal of preclinical disease (Kivisaari, Monsch, & Taylor, 2013; Mortamais et al., 2016). Specifically, feature-based models of semantic memory predict a recognition deficit for objects characterised by numerous shared features (e.g., living objects) vs. relatively few distinct features (e.g., non-living objects) following degeneration of the PRc (Kivisaari, Tyler, Monsch, & Taylor, 2012; Taylor et al., 2011; Taylor, Moss, & Tyler, 2004; Tyler et al., 2004).

The differential demands placed by living vs. non-living objects on PRc function is leveraged by Gallery Game to enhance sensitivity to early cortical tau-related degeneration, as follows. Non-living objects tend to have a distinct form-to-function mapping and hence are easier to discriminate than living objects, which tend to share multiple correlated features (e.g., ‘has eyes’, ‘has ears’, ‘ has four limbs’; (Taylor et al., 2004; Tyler & Moss, 2001). The distinguishing features of living objects (e.g. the hump(s) of a camel) tend to be weakly correlated with the common features, which in turn leads to a more vulnerable representation of these discriminatory characteristics in PRc-dependent semantic memory. For example, confrontation naming of living objects is differentially impaired in a mild AD group, where performance deficits are correlated with PRc atrophy (Kivisaari et al., 2012). Similarly, adults with amnestic mild cognitive impairment (aMCI) or mild AD show greater false recognition (or ‘false alarms’) to novel living objects than non-living objects after implicit learning (Kivisaari, Monsch, et al., 2013). In addition, discrimination of novel items is linked more strongly to PRc function than recognition of familiar objects (supported by ventral parietal regions) (Krumm et al., 2017). Drawing on this collection of evidence, Gallery Game’s test of delayed recognition is designed to provide a sensitive marker of PRc dysfunction – and hence early AD-related pathology – by presenting living and non-living target images amid highly-confusable matched but novel distractors.

Targeting the cognitive processes supported by the ERc and hippocampus alongside PRc-dependent function strengthens the sensitivity of Gallery Game to early Braak pathology in the MTL. The ERc and hippocampus are critical for the integration of contextual information within memory, for example the binding of object and location (Carr et al., 2017; Kivisaari, Probst, & Taylor, 2013; Staresina & Davachi, 2009). This form of associative learning is vulnerable in the very early stages of AD (de Rover et al., 2011; Olson, Page, Moore, Chaterjee & Verfaellie, 2006; Sapkota, van der Linde, Lamichhane, Upadhyaya & Pardhan, 2017) including at short durations considered to still be within working or short-term memory. Gallery Game employs a cross-modal paired-associate learning task to further stretch this system, both during immediate learning trials and across much longer retention intervals.

To assess the validity of Gallery Game, we conducted a proof of concept study with a month-long schedule of daily assessments involving encoding, recognition, and recall. In addition, the task was administered to a sample of mid-age adults with a self-selected bias towards familial AD risk, a highly relevant sample for the detection of preclinical AD (Finch, 2009; Irwin, Sexton, Daniel, Lawlor, & Naci, 2018). We predicted that increasing delays between learning and consecutive recognition and recall tests would expose greater individual differences in correct memory retrieval, supporting the value of ecologically valid, long-term measures of episodic memory to detect subtle behavioural differences in populations at-risk populations. Construct validity was assessed by correlating behavioural outcomes with those from established neuropsychological tests of delayed memory. Furthermore, to provide preliminary insight into the utility of false alarms for living as compared to non-living distractors for the detection of preclinical AD, this profile of recognition errors was compared with established screens for global cognitive impairment.

## 2. Methods

### 2.1 Participants

Descriptive data for 35 adult volunteers (97% Caucasian, aged 40-59 years) are shown in Table 1. These participants were recruited from the Oxford study-site of the PREVENT dementia programme (Ritchie & Ritchie, 2012) which excluded individuals with diagnosable dementia. The PREVENT programme intends to recruit 700 individuals who will be phenotyped longitudinally with the aim of defining the interactions of risk factors for dementia with AD biomarkers in mid-age. Although family history of dementia was not an inclusion criterion, participants appeared to self-select for this characteristic: a high proportion (66%) reported a first-degree relative with dementia (43% AD). APOE4 status is collected but remains blinded until the end of study recruitment (expected by end of 2019).

**Table 1.**
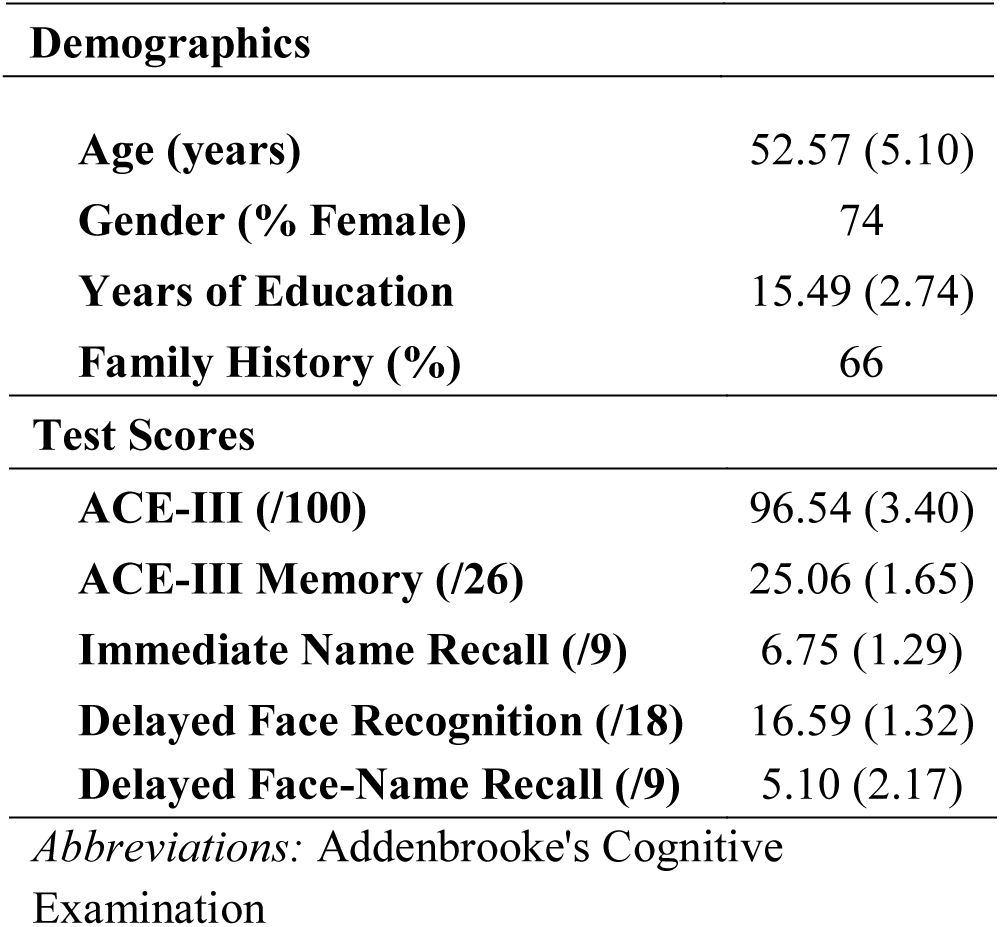
Participant demographics and neuropsychological test scores shown as mean (*SD*) unless stated otherwise.

The current study was ethically approved (University of Oxford Medical Sciences Inter-Divisional Research Ethics Committee: R48717/RE001) and compliant with the Helsinki Declaration of 1975. Written consent was required upon entry to the study.

### 2.2 Neuropsychological Assessment

A battery of neuropsychological tests was completed during a site visit as part of the main PREVENT study (Ritchie & Ritchie, 2012). For the purpose of the current study, the total and episodic memory scores on the Addenbrooke’s Cognitive Examination Revised (ACE III-R) (Noone, 2015) were used as an indicator of general cognitive and episodic memory ability respectively. Episodic memory sub-tests of the computerised COGNITO battery (de Roquefeuil Guilhem, 2014), specifically: 1) immediate name recall (/9), 2) delayed face-name association recall (/9) and 3) delayed face recognition (/18), were selected for comparison with the Gallery Game smartphone task.

### 2.3 Gallery Game

Gallery Game is a freely available cognitive task deployed within the Mezurio smartphone app platform (https://mezur.io). The task has a repeated-measures design, with each participant asked to complete daily assessments consisting of multiple learning tasks, each associated with a single recognition memory and recall test. The current study utilised a 30-day testing phase to ensure this proof-of-concept had sufficient statistical power to interrogate behavioural outcomes, however, future research is not restricted to a 30-day assessment schedule.

#### 2.3.1 Stimuli

Photographic stimuli of living and non-living concepts, isolated on a white background, were included in Gallery Game. Images from the Bank of Standardised Stimuli (BOSS) (*n*=480) (Brodeur, Dionne-Dostie, Montreuil, & Lepage, 2010) were used in the initial administration of Gallery Game, however, subsequent updates to the Mezurio App included a wider selection of professionally photographed stimuli (*n* = 1200) licensed from Shutterstock (https://www.shutterstock.com), an online image platform. Metrics of familiarity, visual complexity and object category were collated for each image stimuli. For the BOSS database, familiarity and visual complexity ratings consisted of human ratings collected along a 5-point Likert scale (Brodeur et al., 2010). The database of Shutterstock images included JPEG compressed file size as a proxy of visual complexity (e.g. Donderi & McFadden, 2005; Machado et al., 2015). Published word frequency between years 1999 and 2009, extracted from Google Ngram (Michel et al., 2011), was used to approximate concept familiarity (Brysbaert et al., 2011). Stimulus category was coded in-house (e.g. Mammals, Vegetables, Clothing), as was the left/right/up orientation of the object image. Stimuli selection was restricted to a single image dataset (BOSS or Shutterstock) for each participant, dependent on when they began their schedule of daily assessments.

#### 2.3.2 Learning task

Gallery Game contains a paired-associate learning task intended to maximally load on entorhinal and hippocampal regions, vulnerable to change in early AD (de Rover et al., 2011; Loewenstein, Curiel, Duara, & Buschke, 2018). An example of a single learning trial is shown in Figure 1a. Each object image was shown along with an arrow pointing left, right, or up in first instance, with participants being required to ‘swipe’ the image in the cued direction to learn the object-direction association. The directional cue was absent on subsequent presentations of this object to test immediate recall for object-direction pairings, with an incorrect response triggering a second object-direction learning trial with the directional cue. Object images were shown in a ‘gallery’, with the number of images presented in each iteration of this gallery growing progressively from 1 to 6 if all object-direction pairings in the preceding iteration were successfully recalled. Within each iteration, images were presented in a random order. After each successful iteration (100% correct), Gallery Game provided positive feedback (a gold star). Each learning task continued until a maximum of 6 images were shown and the individual reached the stopping criterion that 5 out of the last 7 presentations of a given stimulus were responded to correctly^1^. Repeatedly presenting object-direction pairings until the criterion for successful learning was reached promotes equivalent representation in long-term memory across participants, enabling fairer interpretation of forgetting rates (Elliott, Isaac, & Muhlert, 2014).

**Figure 1.**
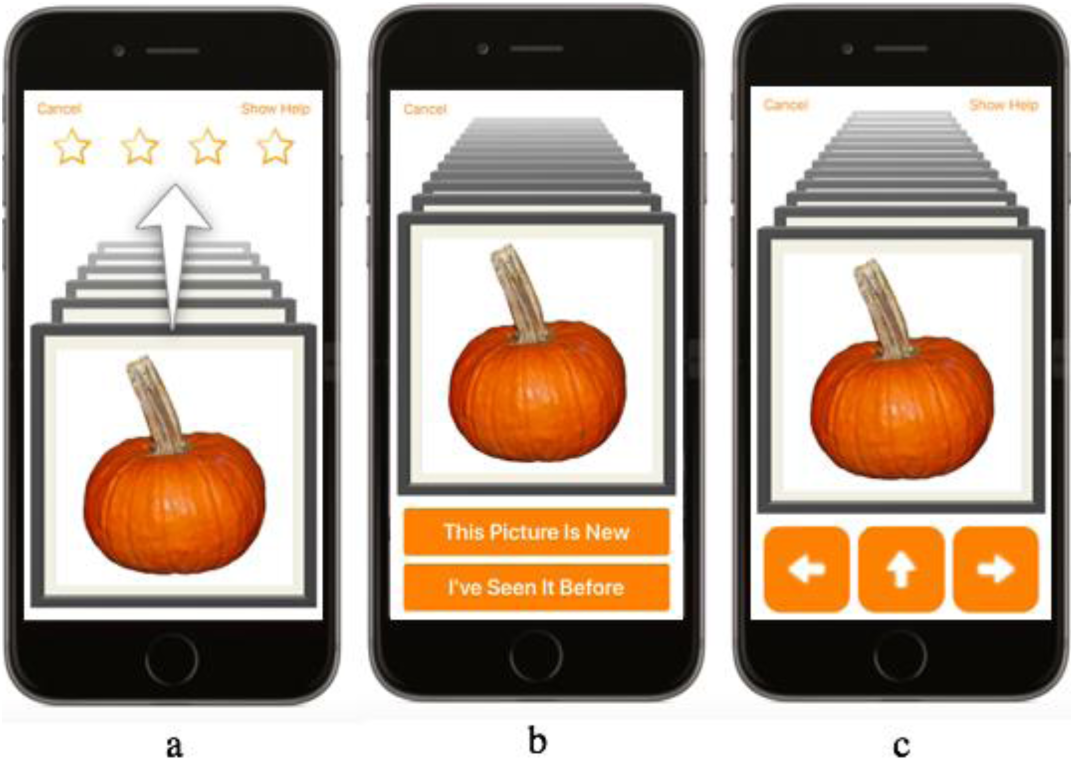
Representation of a learning (a), recognition (b) and recall trial (c) in Gallery Game

Each learning task was scheduled to include up to 6 unique target images (3 living, 3 non-living). Each living target was matched with a non-living target using multivariate genetic matching (visual complexity, familiarity) computed within the ‘Matching’ R package (Diamond & Sekhon, 2012; Sekhon, 2011). Swipe directions (left, right or up) were randomised on a task by task basis, with an addition constraint implemented which biased objects belonging to the same semantic category (e.g. birds, vehicles) being paired with different swipe directions. In addition, the congruency of each image orientation (e.g. the bear facing right) and the swipe direction (left, right or up) was recorded for subsequent analysis.^2^ Across the schedule of daily learning tasks, matched-target pairs were presented in a random order for each participant across all iterations.

#### 2.3.4 Recognition test

A single recognition trial is shown in Figure 1b. In each trial, participants were instructed to classify each image as either a previously seen image or a new image. Each recognition test included 6 target images taken from the same learning task, and 6 matched distractors, presented in a random order. Matching of target-distractor pairings was completed by forming a composite of the visual complexity and familiarity ratings available for each image, and using the Munkres algorithm to produce the assignment solution with the smallest difference cost (Munkres, 1957). Examples of living and non-living target-distractor pairings can be seen in Figure 2.

**Figure 2.**
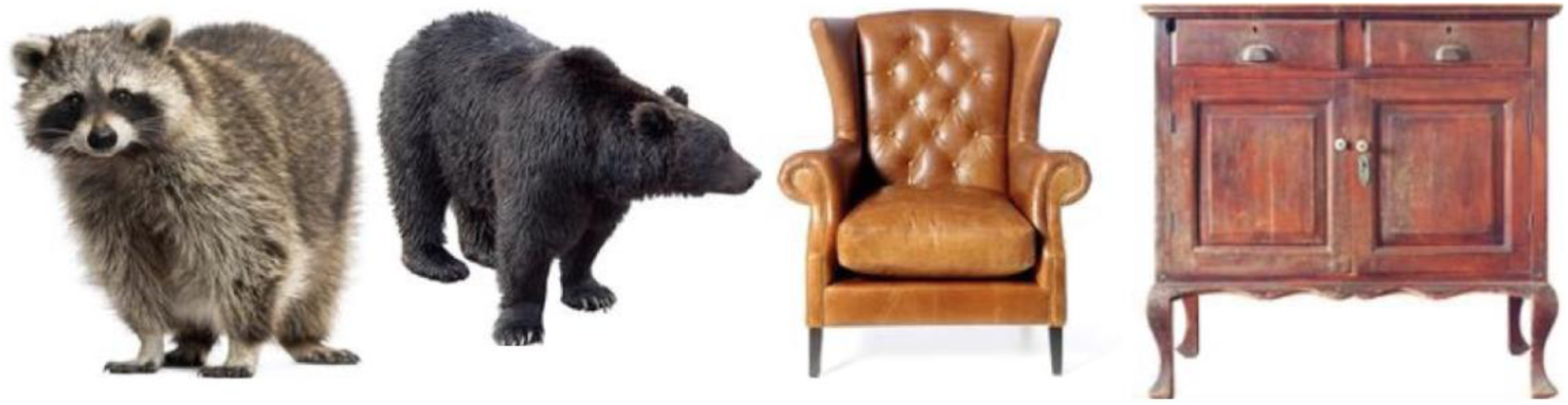
Examples of a matched living target-distractor pairing and non-living target-distractor pairing included in the Gallery Game task.

#### 2.3.5 Recall test

Participants were asked to retrieve the swipe direction (left, right, up) learnt in association with each target image (see Figure 1c for example recall trial). Target images from the same learning task were presented in a random order, with each image presented 3 times per recall test.

### 2.4 Procedure

Participants were invited to download the Mezurio App onto either their own smartphone (Apple or Android) or a loaned Android device. Written instructions and a unique ID were provided by the research team. Participants were prompted to complete daily cognitive tasks within Mezurio for up to 36 days, with the first 30 days including Gallery Game tasks. The app encouraged participants to play at the same time each day by sending a phone-based notification, with a reminder notification sent 15-minutes later if the task was not initiated. Participants had 16 hours to complete each activity after receiving the prompt; tasks not completed within this time window ‘expired’. The schedule adapted to any missed activities to ensure recognition and recall memory was only tested for learnt material. The researchers monitored participant compliance through the ‘Gallery Attendant’ software (freely available as part of the Mezurio platform).

An opportunity to practice the learning, recognition and recall tasks included in Gallery Game was provided at the start of the month’s interactions. During practice, participants were encouraged to use a strategy to learn the association between objects and their ‘swipe’ directions. Example strategies used visual features of the images (e.g. the iron points up) or a semantic storyline (e.g. the wind blows the plant’s leaves to the left’) to assist learning. The decision to introduce strategies at the onset of testing was motivated to reduce between-participant noise in strategy adoption, often related to differences in educational background or cognitive reserve (Harvey, 2017).

Excluding practice days, participants completed up to 22 learning tasks across the month of interactions. For each learnt set of stimuli, participants were prompted to complete consecutive tests of recognition and recall memory after a RI of either: 1, 2, 4, 6, 8, 10, or 13 days. The exact retention interval between learning and test varied as a function of how punctually participants responded to the task notification. Equal numbers of each RI were built into each Gallery Game schedule, split across 3 repeating test blocks across the month. The 13-day RI was introduced into the schedule partway through data collection after a preliminary data analysis showed participants were performing substantially above chance on recognition and recall tests after 10-day delays. As recognition and recall performance after a 13-day RI is only available for a subset of participants (*n*=10), data is excluded from the present statistical analysis.

### 2.5 Statistical analysis

Participants were not asked to emphasise response speed; hence accuracy is considered the primary outcome of Gallery Game performance. Practice trials were excluded prior to analyses, with the exception of a preliminary screen for practice effects on learning task performance. Data was screened for outliers (group mean ± 2 standard deviation (SD)) prior to analysis, with the exception of the ratio of recognition false alarms for living compared to non-living distractor images, as this outcome is intended to provide a signal of very early AD-related deviance in recognition memory outcomes (Kivisaari, Monsch, et al., 2013).

To explore the effect of increasing delays between learning and memory (recognition, recall) test, continuous RIs were grouped as: **1)** 0 > 1.5, **2)** 1.5 > 3, **3)** 3 > 5, **4)** 5 > 7, **5)** 7 > 9, and **6)** 9 > 11 days, with the number of trials completed at each RI shown in Table 2. Results are interpreted in light of both traditional significance testing (*α*=.05) and effect size (η^2^_p_,: small =.01, medium =.06, large=.14, Spearman’s ρ and Pearson’s *r:* small =.2, medium =.5, large =.8) to better understand behavioural outcomes (Ferguson, 2009; Maher, Markey, & Ebert-May, 2013). A Bonferroni-correction is applied for all multiple comparisons unless otherwise noted.

**Table 2.** The number of trials and number of participants contributing data at each RI, shown alongside proportion accuracy of recognition and recall for target images (mean ± SD). *Note:* An equal number of novel matched-distractor images was also presented at recognition test.

| RI (days) | Recognition |  |  | Recall |  |  |
| --- | --- | --- | --- | --- | --- | --- |
|  | Trials (*n*) | Participants (*n*) | Accuracy | Trials (*n*) | Participants (*n*) | Accuracy |
| 0 - 1.5 | 438 | 27 | .92 ± .15 | 1350 | 27 | .77 ± .33 |
| 1.5 - 3 | 424 | 31 | .92 ± .15 | 1326 | 31 | .76 ± .33 |
| 3 - 5 | 504 | 34 | .84 ± .16 | 1494 | 34 | .63 ± .38 |
| 5 - 7 | 479 | 33 | .80 ± .22 | 1437 | 33 | .62 ± .38 |
| 7 - 9 | 462 | 32 | .71 ± .25 | 1404 | 31 | .58 ± .38 |
| 9 - 11 | 474 | 33 | .67 ± .23 | 1458 | 33 | .49 ± .38 |

#### 2.5.1 Learning task

The average number of errors per learning task was treated as the primary outcome for immediate recall. As participants are prompted to repeat the preceding learning iteration following an error, with the number of repeated trials varying from 1 – 6 depending on the size of the iteration when the error is made, proportion accuracy is not selected as an outcome. Due to the progressive increase in object-direction paired-associates, errors are expected to be consistently low.

The association of practice, proxied as the number of learning tasks previously completed, with immediate recall errors was explored using a linear regression analysis. Practice effects were predicted to be minimal given that participants were presented with a learning strategy during practice, and given the unique set of stimuli included in each task. In addition, the effect of image dataset (BOSS, Shutterstock) on immediate recall errors was tested using a Mann-Whitney *U* test as a preliminary check for differences in memorability. Immediate recall errors were log-transformed to limit the effect of positive skew, and then included in a 4 (Congruency of image orientation and swipe direction: neutral, congruent, incongruent – opposite, incongruent – perpendicular) × 2 (Animacy: living, non-living) repeated-measures analysis of variance (ANOVA) to further test stimuli effects.

Galley Game immediate recall errors were correlated with COGNITO immediate name recall, delayed face-name paired-associate recall, and the memory sub-score of the ACE-III-R to assess construct validity.

#### 2.5.2 Recognition test

Performance on individual recognition trials are classified as: 1) Hits -correct recognition of a previously seen targets, 2) Misses-failure to recognise previously seen targets, 3) False alarms – incorrect recognition of a novel distractors, and 4) Correct rejections – novel distractors marked as ‘new’. Hits as a proportion of recognition trials including previously seen targets were included in a 6 (RI: 0 > 1.5, 1.5 > 3, 3 > 5, 5 > 7, 7 > 9, 9 > 11 days) × 2 (Animacy: living, non-living) ANOVA. Pairwise comparisons were used to test both at which point there was a significant change from accuracy at the shortest RI, and if subsequent increases in RI were associated with a significant further decrease in accuracy. Recognition accuracy (hits + correct rejections/total trials) was extracted for correlation with COGNITO face recognition accuracy, however, this only provides a partial account of recognition performance in forced-choice paradigms (Stanislaw & Todorov, 1999). A more complete metric is the *d*’ sensitivity statistic (signal detection theory) (Macmillan & Creelman, 2005), which accounts for the probability of hits as a function of false alarms (Pallier, 2002), computed for the present dataset with a log-linear correction to account for ceiling or floor performance (Brown & White, 2005; Hautus, 1995). In addition, response bias (*b*) was calculated (No bias: *b*=1, bias towards marking images as previously seen: *b* < 1, bias towards marking images as new: *b* > 1). Separate *d*’ and *b* statistics were calculated for living and non-living images, however, these metrics were not extracted for each RI as the number of data points available does not support the normality assumption underpinning signal detection theory.

The proportion of false alarms to living versus non-living distractors, with scores less that 0 representing a higher proportion of false alarms to living distractors, was correlated with the ACE-III-R overall score and memory sub-score.

#### 2.5.3 Recall test

Recall accuracy was subject to a 6 (RI) × 2 (Animacy) × 4 (Congruency) repeated measures ANOVA. Overall recall accuracy was correlated with delayed name-face paired associate recall scores (COGNITO), in addition to the memory sub-score of the ACE-III.

## 3. Results

### 3.1 Learning task

The number of learning tasks previously completed, included as a marker of practice effects, was not significantly associated with variance in the number of immediate recall errors in subsequent learning tasks (*R^2^* =.00, *p*=.453). Following the exclusion of practice trials, participants learnt an average of 99.63 unique target images (*SD*=20.67, range=6-132). Across participants the mean number of errors made per daily learning task was 1.63 ± 2.14, with one participant classified as an outlier (*M*=10.61), hence excluded from further analysis of immediate recall performance.

There was no significant difference in the number of immediate recall errors per learning task for images selected from the BOSS (*M*=1.82) compared to the Shutterstock database (*M*=1.44) (*p*=.305). The main effect of object orientation– direction congruency on log-transformed immediate recall errors approached significance, *F*(1, 3)=2.59, *p*=.054, η^2^_p_ =.029, driven by a marginal difference in errors for congruent (M=.23) and incongruent -perpendicular (*M*=.41) object-direction associations (*p*=.009, Bonferroni corrected α=.008). All other group comparisons, including those between neutral (*M*=.26) and incongruent-opposite (*M*=.26) pairings were non-significant (*p*>.008). In addition, the main effect of animacy (*p=*.889, η^2^_p_ =.000) and Congruency × Animacy interaction were both non-significant (*p=*.675, η^2^_p_ =.006).

Correlations between Gallery Game immediate recall errors and both delayed paired-associate face – name recall (COGNITO), *ρ*(33)=-.53, *p<*.001, and ACE-III memory sub-score, *ρ*(33)=-.44, *p=*.009, were significant (medium effect size). Although non-significant, the correlation between Gallery Game immediate recall errors and delayed name recall (COGNITO) was in the anticipated direction, *ρ*(33)=-.20 *p*=.258.

### 3.2 Recognition test

An average of 171.88 recognition trials was completed by each participant across the period of Gallery Game assessment (*SD*=52.57, range: 72-264), with the distribution of data points available for each RI shown in Table 2. One participant, who only learnt 6 images, did not complete any recognition or recall trials and hence is absent from ongoing analyses.

#### 3.2.1 Recognition accuracy

The main effect of RI on recognition of previously seen targets was large, *F*(5, 365)=17.20, *p*>001, η^2^_p_ =.191 (see Figure 3). Accuracy was high following the shortest RI (<1.5 days) (*M*=.93 ± .15), with no significant decline in performance at RIs of 1.5 < 3 days (*M*=.92 ± .15, *p*=.975) or 3 < 5 days (*M*=.84 ± .16, *p*=.030). There was a significant decrease in accuracy of target recognition following a 5-7 days RI (*M*=.80 ± .22, *p*=.002). In comparison, there was a marginal stepwise decline in accuracy for RIs of 7-9 days (*M*=.71 ± .25, *p*=.012), but the decline in performance was not significant for a further increase (*M*=.67 ± .23, *p*=216). Longer RIs are also associated with greater variation in performance between individuals. Recognition accuracy was significantly higher for living (.85) than non-living stimuli (.77), *F*(1, 365)=15.35, *p*<.001, η^2^_p_ =.040), however, the RI × Stimuli Type interaction was non-significant (*p*=.847, η^2^_p_ =.004). Total recognition accuracy (targets, distractors) was non-significantly correlated with COGNITO face-name recognition, *r*(32) =.25, *p*=.152, however, the association was small and in the anticipated direction.

**Figure 3.**
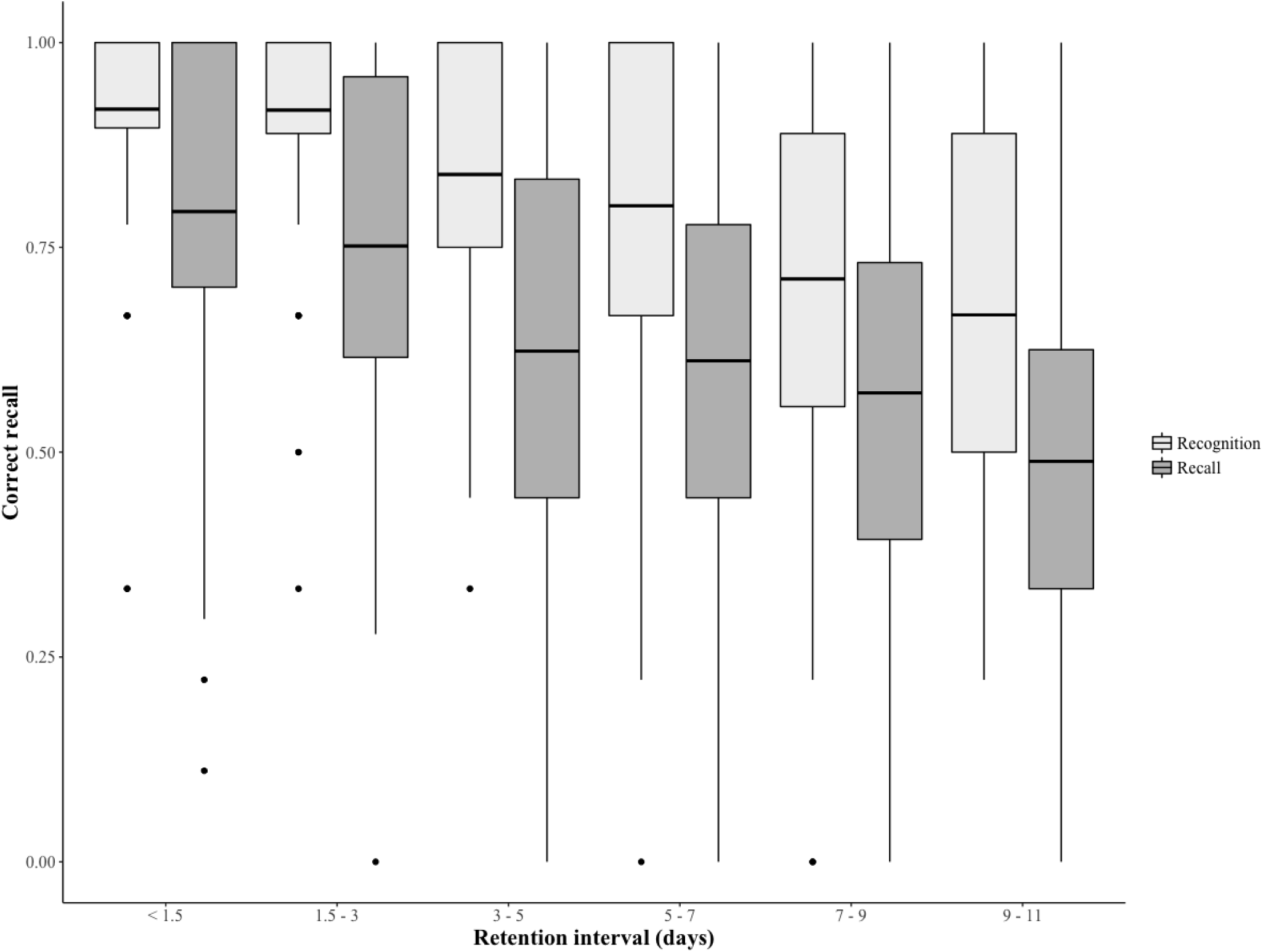
Accuracy (proportion) of target recognition and target object -direction paired-associations recall at each RI.

#### 3.2.2 Recognition sensitivity

Recognition sensitivity (*d*’) was normally distributed (Shapiro-W=.977, *p*=.674, *M*=2.11 ± 1.18), with a mean *b* of 1.59 ± 1.18. There was no significant difference in *d*’ for living (*M*=2.19 ± .87) and non-living objects (*M*=2.12 ± .70), *t*(33)=.44, *p=*.664. Overall *d*’ did not significantly correlate with ACE-III-R memory scores (*ρ*=.36, *p*=.034) or COGNTIO face recognition *(r*=.24, *p*=.171) (α=.025), however the associations were small and in the anticipated direction with both established tests.

#### 3.2.3 False alarms

Non-living distractors (.898) were associated with a higher proportion of correct rejections than living distractors (.835), *t*(33) = −3.59, *p*=.001. Two participants were classed as outliers for the proportion of living compared to non-living false alarms (*M*=.93 ±.12). Although the correlation between this ratio and ACE-III-R score was non-significant (ρ=.06, *p*=.734) (see Figure 4), these two participants also demonstrated outlier performance on the ACE-III-R (*M*=96.17±3.37), with one participant scoring below the recommended cut-off (88/100) for cognitive impairment (Mioshi, Dawson, Mitchell, Arnold, & Hodges, 2006). The correlation between the false alarm ratio and ACE-III-R memory score was non-significant (*ρ*=.08, *p*=.671)(but note negative skew in ACE-III-R memory scores), however, construct validity of the signal of false alarms for living versus non-living stimuli is supported by a positive correlation with COGNITO face recognition scores (*r*(33)=.54, *p*<.001).

**Figure 4.**
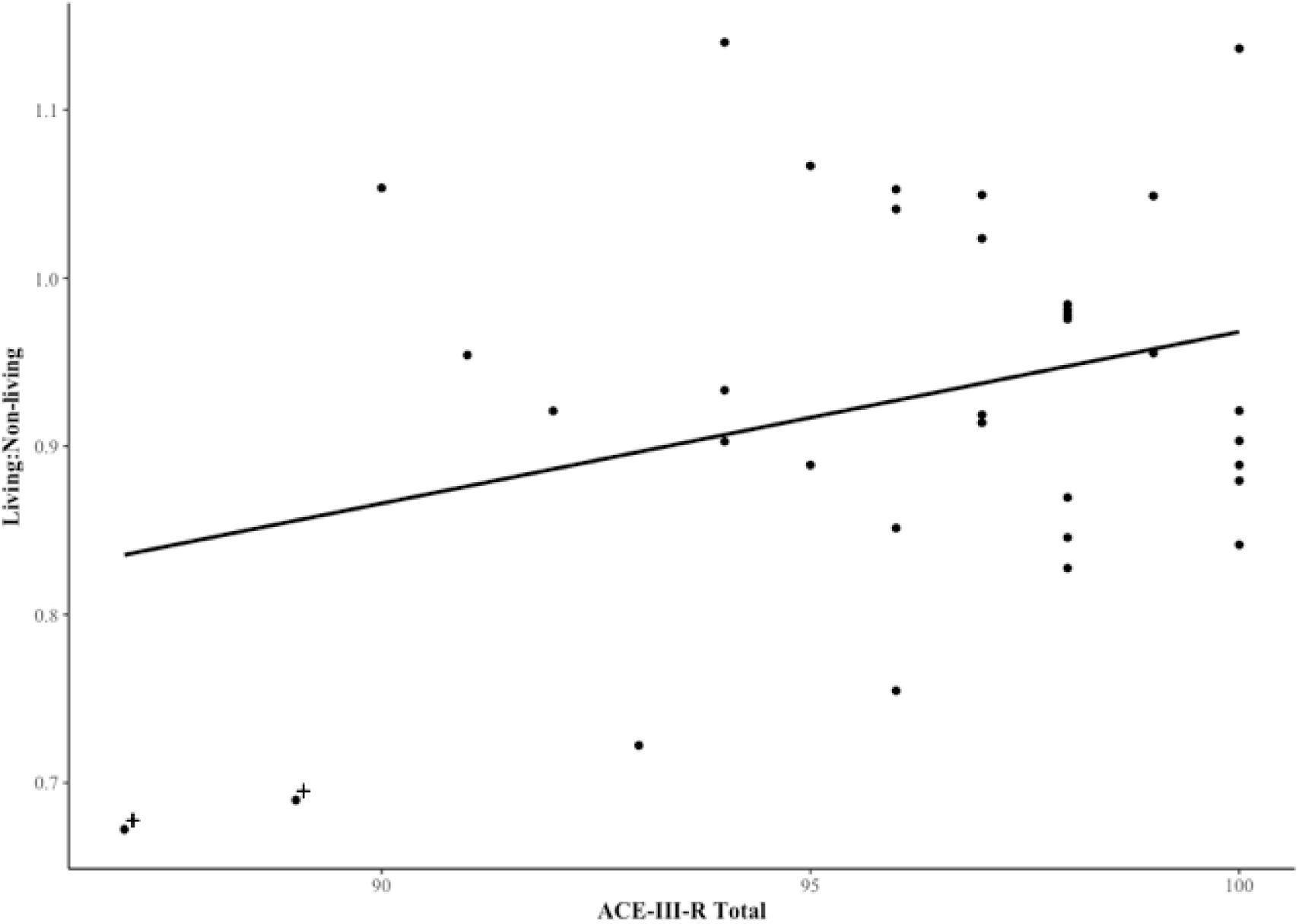
Correlation between the ratio of false alarms for living versus non-living distractors and ACE-III-R scores. *Note:* Outliers are denoted by a +

### 3.3 Recall test

An average of 260.74 recall trials was completed by each participant across the period of Gallery Game assessment (*SD*=81.90, range: 108-396), with the distribution of data points available for each RI shown in Table 2.

The main effect of RI on object-direction paired-associate recall was significant, *F*(5, 1205)=20.43, *p*<001, η^2^_p_ =.072, driven by a consistent decrease in accuracy with increasing RIs up to 11 days (see Figure 3). The decrease in recall accuracy was non-significant with an increase in RI from <1.5 days (*M*=.77±.33) to 1.5 > 3 days (*M*=.76 ±.33)(*p*=.707), however, recall performance was significantly reduced after a RI of 3 > 5 days (*M*=.63±.38)(*p*<.001). Accuracy at retention intervals 3 > 5, 5 > 7 (*M*=.62±.38) and 7 > 9 (*M*=.58±.38) days was not significantly different (*p*>.01), however, there was a further significant drop in recall performance following a delay of 9 > 11 days (*M*=.47, *p*=.38) (*p*=.001). There was a significant main effect of congruency, *F*(3, 17.69)=17.69, *p*<001, η^2^_p_ =.04, driven by recall of object – direction pairings being more accuracy for congruent associations (*M=*.75±.33) than neutral (*M*=.64±.38), incongruent – perpendicular (*M*=.55±.37) and incongruent – opposite (*M*=.61±.41) pairings (*p*<.001). Accuracy for neutral pairings was also significantly greater than accuracy for incongruent -perpendicular pairings (*p*=.002), with all other pairwise comparisons non-significant (*p*>.008). The main effect of animacy and all interaction terms were non-significant (*p*>.05).

There was a significant positive relationship between overall recall accuracy for image-swipe directions in Gallery Game and delayed face-name association recall in the COGNITO battery, *r*(31)=.40, *p*=.022. The relationship between Gallery Game recall and ACE-III memory sub-scores was in the predicted direction, *r*(31)=.34, *p*=.063, but failed to reach significance. This may in part be due to 58% of participants being at ceiling on this task.

## 4. Discussion

This proof of principle study explored Gallery Game performance in a sample of mid-age adults with a high proportion at elevated familial risk for AD. The results confirmed the validity of the design of this novel smartphone assessment, with clear associations between its outcomes and those from established in-clinic neuropsychological tests of episodic memory.

Specifically, immediate recall accuracy in the current sample was predominantly at ceiling across learning tasks, highlighting the value of progressive, repeated learning iterations to promote equivalent encoding of object – direction paired-associates between participants. Variation in both object recognition and free recall performance supported the utility of Gallery Game for detecting episodic memory differences in adults with no clinical cognitive impairment. As anticipated, increasing the RI between learning and test was associated with a medium-large monotonic decrease in memory retrieval. As expected, forgetting was non-linear with increasing delays. Furthermore, the current data confirms the construct validity of Gallery Game outcomes (immediate and delayed-recall, delayed recognition) by demonstrating the predicted associations with highly relevant paired-associated memory measures from both COGNITO (de Roquefeuil Guilhem, 2014) and the memory sub-score of ACE-III-R (Noone, 2015). Consideration of the proportion of false alarms for living compared to non-living distractors in relation to stringent diagnostic cut-offs for global tests of cognitive impairment (ACE-III-R) (Mioshi et al., 2006) provides preliminary support for the utility of this behavioural marker as a signal of very early cognitive impairment.

Endpoints derived from Gallery Game were comparable to ‘gold-standard’ in-clinic episodic memory assessments, with this confirmation of construct validity representing a critical step towards establishing the task’s utility as a clinical tool (Chinner et al., 2018; Jongstra et al., 2017; Moore, Swendsen, & Depp, 2017). Whilst many available digital tools are a direct translation of existing pen-and-paper neuropsychological tests, Gallery Game measures trial-by-trial performance in a paradigm specifically designed for high-frequency, longitudinal smartphone assessment; essential for monitoring the progressive change in behaviour characteristic of AD (Onnela & Rauch, 2016). There was no evidence of practice effects across this month-long schedule of daily learning tasks, promising for Gallery Game’s sensitivity for capturing longitudinal decline. Further work, however, is required to identify which simple behavioural outcomes from this task are most sensitive to the functional-neuroanatomical changes characteristic of early AD-related degeneration, for example by correlating recognition and free recall scores with anterior MTL regions of interest scores based on magnetic resonance imaging scans, and hence of greatest predictive utility.

The current data supports the value of extending delays between learning and memory retrieval to increase measurement sensitivity to performance differences in healthy adults, with accuracy of object recognition in particular consistently high at shorter RIs. In further support of the importance of longer-term RIs, neuropsychological tests conforming to standard memory delays of approximately 30 minutes (Lezak et al., 2012) show negative skew in the present sample (specifically the ACE-III-R memory sub-score and COGNITO face recognition). One key strength of remote digital assessment is the ability to frequently sample memory retrieval across diverse, variable RIs, and the current study provides the first repeated-measures interrogation of long-term memory across more than five RIs (Fisher & Radvansky, 2018). By contrast, participant burden has restricted the bulk of prior research to an immediate, delayed (30 minute), and long-term memory test (e.g. van der Werf et al., 2016; Zimmermann & Butler, 2018).

By including a greater sample of RIs, Gallery Game enables further interrogation of the form of forgetting (Averell & Heathcote, 2011), represented as a non-linear function defined by a period of enhanced forgetting followed by a more subtle loss of information from memory. This initial dataset shows a significant drop in free-recall performance after three days and in object recognition after five days, with second declines in memory performance at RIs seven - nine and nine – 11 days respectively. Recent research suggests a transition in memory retention after a delay of seven days (Fisher & Radvansky, 2018), accounted for by distinct, time-dependent consolidation processes. Further study of this shift in memory retention will allow for more efficient Gallery Game administration, targeted around this potentially sensitive forgetting period. In addition, ongoing work utilising the Gallery Game task will examine between-participant group differences in the form of forgetting, specifically whether a consistent trajectory of accelerated forgetting or divergence following an identified delay is most predictive of cognitive impairment (Averell & Heathcote, 2011; Castel, Balota, Hutchison, Logan, & Yap, 2007).

Increased confusability of living compared to non-living objects as a hypothesised manifestation of perirhinal neurodegeneration is central to the utility of Gallery Game as a marker of preclinical AD. Of interest, although participants were screened for cognitive impairment at recruitment, differences in the proportion of false alarms for novel living compared to non-living distractors correlated with outlier performance on the ACE-III-R, perhaps consistent with the idea that these individuals are showing very early cognitive impairment specific to PRc function (Mioshi et al., 2006). Although such conclusions are necessarily speculative at the present sample size, this finding is promising for future use of Gallery Game in the detection of preclinical AD. Future work should seek to validate this prediction based on Gallery Game endpoints against cerebrospinal fluid, positron emission tomography (PET) and magnetic resonance biomarkers indicative of AD-pathology and subsequent cognitive impairment.

Although this research is a critical first step in establishing the scientific value of Gallery Game ahead of more costly clinical validation work and deployment in large-scale cohorts, there are limitations worthy of note. Statistical significance of group-level effects must be interpreted with caution, chiefly as variable participant adherence leads to non-uniform sampling of memory retrieval at each retention interval. This is an expected risk of self-administered, daily cognitive assessment (e.g. Allard et al., 2014; Schweitzer et al., 2017). The present sample size limits well-powered interrogation of within-participant variation and stimuli-level effects, for example linked to familiarity or emotional valence (Khosla, Raju, Torralba, & Oliva, 2015). In addition, the sample consisted of individuals with high levels of education which may influence generalisability of the current findings. An ongoing deployment in a large e-cohort will aim to address the issues of sample size and generalisability, including granular analysis of individual-level performance, essential for maximising the value of digital measurement in clinical research (Resnick & Lathan, 2016).

### 4.1 Conclusions

This proof of concept study provides a preliminary validation of the Gallery Game, a newly developed smartphone task, as a self-administered measure of long-term episodic memory suitable for use in non-clinical populations. Furthermore, the increase in variance captured by increasing the length of delay between learning, recognition and recall test suggests including ecologically relevant tests of true long-term memory may facilitate detection of AD at an earlier stage of progression. Preliminary exploration of the semantic processing deficits hypothesised to signal the very earliest emergence of tau pathology are promising for the utility of Gallery Game as a screen for preclinical AD, justifying the ongoing validation of this task against biomarkers of degeneration.

## Disclosure Statement

Chris Hinds is funded by the Robertson Foundation and acknowledges support from the NIHR Oxford Health Biomedical Research Centre, a partnership between Oxford Health NHS Foundation Trust and the University of Oxford. Claire Lancaster is jointly funded by Eli-Lilly and F. Hoffmann-La Roche Ltd. Chris Chatham and Kirsten Taylor are full-time employees of F. Hoffmann-La Roche Ltd. The PREVENT research programme was developed with a grant from Alzheimer’s Society. Ivan Koychev is funded through the National Institute of Health Research Integrated Academic Training scheme (CL-2015-13-007) and declares support for this study through Oxford Clinical Academic Graduate School Clinical Lecturer Support Scheme, the Academy of Medical Sciences Clinical Lecturer Starter Grant (SGL016\1079) and the Medical Research Council Deep and Frequent Phenotyping study grant (MR/N029941/1). Amy Chinner and Jasmine Blane are funded in part through the Oxford Health Biomedical Research Centre.

## Footnotes

1 Subsequent to this first study, a second learning criterion has been implemented within Gallery Game to limit the number of trials available if a participant consistently falls below 100% accuracy on learning iterations. Participants may attempt each size of learning iteration (up to 6 images) 10 times, with a maximum of 128 stimuli presentations per learning task.

2 Subsequent to this first study, image category, orientation and swipe direction are now balanced within each individual learning task.

